# Transcriptomic responses to *Marteilia sydneyi* infection in the Sydney rock oyster *Saccostrea glomerata*

**DOI:** 10.1101/2025.02.02.636094

**Authors:** Nikolina Nenadic, Ido Bar, Carmel McDougall

## Abstract

*Marteilia sydneyi*, an ascetosporean parasite, is the causative agent of Queensland Unknown (QX) disease in *Saccostrea glomerata*. QX disease outbreaks often lead to high mortality rates and considerable population losses. Investigating host/parasite interactions at a molecular level is imperative to better understand how *S. glomerata* mounts immune defences, and to explore whether *M. sydneyi* evades host responses. This study aims to investigate *S. glomerata’s* response to *M. sydneyi* infection through differential gene expression analysis to uncover immune mechanisms and potential markers for resistance. RNA sequencing and differential gene expression analysis revealed widespread transcriptional changes between infected and non-infected oysters. Genes encoding proteins involved in pathogen recognition and immune response signalling, such as galectin-4-like and G-protein coupled receptors, were significantly differentially expressed in *S. glomerata* infected with *M. sydneyi*, suggesting involvement in the host’s immune responses. Moreover, the upregulation of cytochrome P450 family genes indicate increases in the host’s detoxification processes and metabolic pathways, a possible response to infection-induced stress. However, extracellular superoxide dismutase, a gene previously implicated in the oxidative stress response to pathogens, was significantly downregulated, suggesting potential suppression of oxidative burst defence mechanisms. These results reveal the complex nature of *S. glomerata’s* response to *M. sydneyi* infection, and possible suppression or evasion of host defences by the parasite. The study also identifies multiple genes that likely play crucial roles in the molecular responses and defence mechanisms of *S. glomerata* to *M. sydneyi* infection. The identification of these genes provides potential target genes for future studies and possible biomarkers for breeding QX*-*resistant oyster lines.

**Highlights:**

- *M. sydneyi* infection induces a large transcriptional response in *S. glomerata*
- Upregulation of pathogen recognition genes, including galectins, occurs in infected tissues.
- Lack of differential expression of apoptosis genes suggests other immune strategies.
- G-protein coupled receptor pathways were upregulated, indicating species-specific responses.
- Results suggest possible suppression of host immune responses by the parasite.

## 1. Background

Oysters play a vital role in both marine ecosystems and global aquaculture, contributing significantly to coastal economies and food production [1–4]. However, the sustainability of oyster aquaculture is increasingly threatened by recurrent outbreaks of parasite-induced diseases [5]. There are a range of different parasites that infect oysters from several different taxonomic groups. A group of prevalent oyster parasites with substantial impact on oyster populations is Ascetosporea, which includes *Mikrocytos mackini* (parasitic in *Magallana gigas, Crassostrea virginica, Ostrea edulis* and *Ostrea lurida*), *Haplosporidium nelsoni* (parasitic in *C. virginica*), *Marteilia refringens,* and *Bonamia ostreae (both primarily parasitic in O. edulis)* [6, 7, 9–11]

Like all other invertebrates, oysters lack an antibody-based adaptive immune system and rely on the innate immune system for defence against pathogens [12, 13]. The innate immune system in oysters relies on specific receptors that are able to recognise broad pathogen-associated molecular patterns (PAMPs) and activate humoral and cellular responses [12, 14]. The humoral response involves the release of antimicrobial proteins and enzymes into the hemolymph where cellular responses are mediated by hemocytes (the primary defence cells in oysters) at the source of infection [15, 16]. Hemocytes are involved in several innate immune functions such as encapsulation, the production of reactive oxygen species (ROS), and phagocytosis [14].

Molecular studies have shown that parasite infections trigger a range of innate immune responses in oysters, often leading to changes in the expression of genes involved in oxidative stress regulation, immune signalling, and pathogen recognition [15, 17, 18] [13,17,18]. For example, genes encoding antioxidant enzymes, protease inhibitors, and pattern recognition proteins have been associated with response to infection and immune regulation [12, 13, 15]. Despite advances in understanding oyster immunity, knowledge gaps remain regarding whether responses to parasites differ from bacterial or viral infections, and the extent of cross-species similarities in immune strategies. Addressing these gaps is essential for improving oyster disease management and selective breeding efforts.

Oyster aquaculture has been a vital industry in Australia since European settlement, with *Saccostrea glomerata* (the Sydney rock oyster) being a key species [21], currently contributing AUD $102 million annually to the national economy [22]. *S. glomerata* populations have suffered significant declines due to a parasite-induced disease, Queensland Unknown (QX) [23]. Since the 1960s QX disease has been responsible for severe losses in wild and farmed *S. glomerata* populations, with up to 95% overall mortality [24], and many consider QX to be the biggest threat to the *S. glomerata* wild populations and aquaculture industry [23, 25, 26]. QX is caused by the protozoan parasite *Marteilia sydneyi*, which enters the oyster through its gills and labial palps, systematically spreading throughout the oyster until it establishes itself within the digestive tubule epithelium [27]. It is there that the sporonts contained in the established sporangiosorae undergo further proliferation, starving and killing the host [28]. In response to significant mortality the industry has turned to selective breeding for disease resistance [29], and it was found that QX-resistant bred *S. glomerata* exhibited stronger immune responses against QX infection compared to wild-type oysters [19].

Previous studies on *S. glomerata* have identified key molecular responses associated with resistance to QX disease, providing insights into the genetic basis of immunity. QX-resistant oysters have demonstrated increased constitutive phenoloxidase activity in their hemolymph and hemocytes, along with larger and more active immune cells [19]. These findings suggest that elevated phenoloxidase activity and a robust phagocytic response play a vital role in defence against *M. sydneyi* infection. Additionally, gene expression analyses have revealed differential baseline levels of key immune-related genes in disease-resistant oysters compared to wild populations. Notably, extracellular superoxide dismutase (EcSOD) exhibits higher baseline expression in resistant oysters, while peroxiredoxin 6 (Prx6) and interferon-inhibiting cytokine factor (IK) showed lower expression levels [30]. These expression patterns are hypothesized to enhance the production of hydrogen peroxide during respiratory bursts, which is crucial for neutralizing pathogens [30].

Despite the economic and ecological significance of QX disease to the Australian oyster industry, the molecular responses of *S. glomerata* to *M. sydneyi* infection remain poorly understood, likely due to the inability to culture the parasite or perform controlled challenge experiments. In contrast, oyster responses to parasite infection in northern hemisphere species have been better studied (e.g., *O. edulis and M. gigas* responses to the rhizarian parasite *B. ostreae*), which has led to the identification of key immune genes and pathways involved in host defence [17, 18, 31, 32]. This knowledge gap underscores the importance of investigating the host responses of *S. glomerata* to *M. sydneyi* during QX infection.

In this study we conducted a comprehensive transcriptomic analysis to investigate the molecular responses of *S. glomerata* to *M. sydneyi* infection. Using RNA sequencing, differential gene expression analysis, and functional annotation, we identified key genes and pathways involved in the oyster’s immune response to QX disease. By comparing gene expression profiles between infected and uninfected oysters, we aimed to gain insights into the molecular strategies employed by the host to combat QX disease.

## 2. Materials and Methods

### 2.1 Sample collection

20 *S. glomerata* oysters (sourced from wild-caught spat) were collected from the Pimpama River, Queensland (Fig. 1) during a QX outbreak event. Oysters were approximately 2.5 years old and had been in the held in the river for a minimum of 3 months. Once collected oysters were held in an aerated aquarium containing natural seawater from the oyster lease at 25°C for 3 days prior to processing and were fed approximately 1 mL of Shellfish Diet 1800™ (Reed Mariculture) daily.

**Fig. 1.**
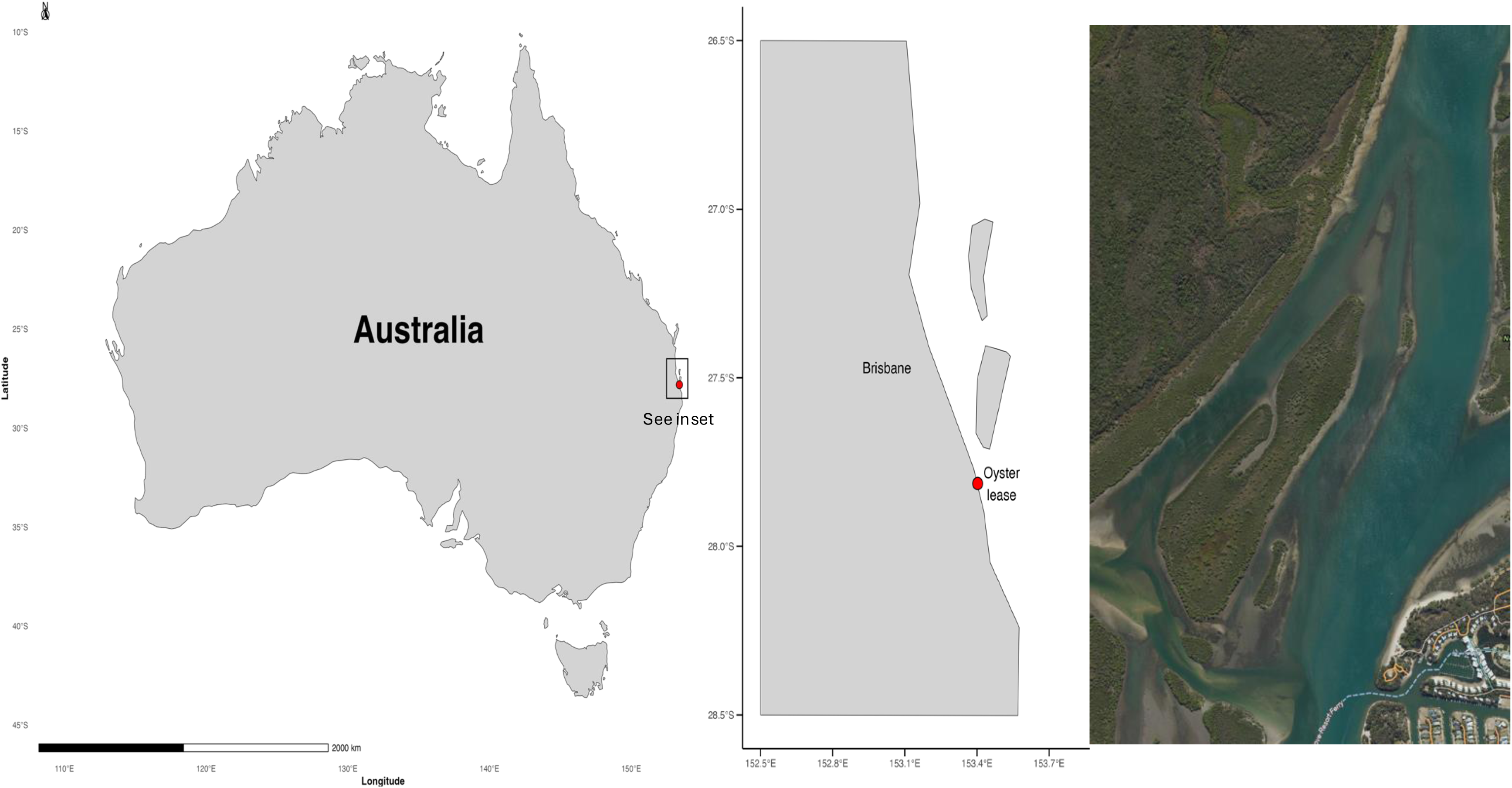
Oyster collection site located on the western side of Woogoompah Island, - 27.813664, 153.402702, in Moreton Bay, QLD. Satellite image © Queensland Globe, State of Queensland, CC BY 4.0.

### 2.2 Oyster sampling and DNA extractions

Each oyster was shucked and the digestive tissue was dissected and split into three samples. The first piece was stored in RNAlater (Sigma) at 4°C for 24 hours and then at -20°C for long-term storage. The second piece was stored in 80% ethanol at 4°C for DNA extractions, and the third was immediately stored at -80°C for subsequent histological examinations.

DNA extractions were performed from ethanol-stored samples using the Qiagen DNeasy Blood and Tissue Kit™ following the protocol provided. A variation was made to the protocol; 30 µL of DNAse/RNAse free water was used in the final DNA elution step instead of the suggested 200 µL of Buffer AE. DNA yield was assessed using the Qubit Fluorometer and the DNA broad range kit (Thermo Fisher Scientific), following the provided protocol. The extracted DNA was diluted to 50 ng/µL with nuclease-free water.

### 2.3 Diagnosis of infection

#### 2.3.1 PCR

Samples were assessed for presence of *M. sydneyi* infection using the Leg1 and Pro2 primers that yield an expected product size of 195 bp [33]. PCR amplifications were performed in 20 µL reactions containing: 2 µL of 10x ThermoPol 10X buffer, 1 µL of each primer at 10 µM, 0.4 µL dNTP solution mix at 10 mM, 0.2 µL Taq DNA polymerase (all PCR components from New England Biolabs), and 1 µL of sample DNA at 50 ng/µL. The following PCR thermoprofile was run on a thermal cycler: 94°C for 2 minutes followed by 30 cycles of 94°C for 30 seconds, 55°C for 30 seconds and 68°C for 1 minute with a final extension of 68°C for 10 minutes and incubation at 14°C. PCR amplicons were visualised on a 2.5% TAE agarose gel.

#### 2.3.2 Histology

Histological screening was performed to confirm infection in samples that tested positive for *M. sydneyi* infection via PCR. The protocol outlined in Adlard and Worthington Wilmer [34] was followed with the variation of using digestive tissue that was frozen at -80°C then thawed prior to smearing. Initial tests showed that this provided identical results to the analysis of fresh tissue (Supplementary Fig. 1). Thawed tissue was blotted on microscope slides and left to dry for 5 minutes. The Hemacolour Rapid Staining of Blood smear kit (Sigma-Aldrich) was used; slides were submerged in solution 1 for 5 seconds, solution 2 for 3 seconds, solution 3 for 2 seconds and solution 4 for 20 seconds, with hand mixing. Slides were left for up to 15 minutes to dry, covered with 100 µL of DPX mounting solution and a coverslip, and left in a dark space to set for 24 hours. Slides were inspected under a microscope for presence of *M. sydneyi* cells.

### 2.4 RNA Extraction and Sequencing

Three samples highly infected with *M. sydneyi* and three non-infected samples were chosen for transcriptome sequencing. Samples were chosen based on PCR amplicon brightness when visualised on an agarose gel and number of sporonts visible upon histological examination. RNA was extracted from 0.5 cm^2^ of the excised digestive tissue stored in RNAlater at -20°C. Extractions were carried out using the standard protocol for RNA extraction using Trizol™ with the following modifications: 200 µL of Trizol™ was added to digestive tissue. The samples were incubated at 65°C for five minutes with vortexing every 2 minutes, the remaining 400 µL of Trizol™ was then added and vortexed for 15 seconds. 20 µL of 1-bromo-3-chloropropane was added to initiate phase separation. For precipitation, 0.5 µL of 20 mg/mL glycogen was added to the transferred supernatant and mixed before the addition of 100 µL of ice-cold isopropanol. The resulting pellet was resuspended in 6 µL of nuclease-free water. RNA yield was assessed using the Qubit Fluorometer RNA broad-range kit following the manufacturer’s instructions. RNA quality was assessed by electrophoresis on a 1% TBE gel. Extracted RNA was sent to Macrogen, Korea for quality control (via Tapestation), library preparation and transcriptome sequencing. Individual libraries were created for each sample using the Truseq stranded mRNA kit (Illumina) and were sequenced on a NovaSeq 6000, generating 150 bp paired-end reads.

### 2.5 RNA-Seq Data Analysis

#### 2.5.1 De Novo Transcriptome Assembly

Paired-end RNA-Seq reads were error-corrected with Rcorrector (v1.5.0, [40]) and any unfixable reads were removed. A de novo transcriptome was assembled and annotated from the error-corrected reads with TransPi (v1.1.0, [41]), a pipeline implemented in the Nextflow scientific workflow system [42]. In brief, TransPi performs a complete transcriptome assembly, basic annotation and assessment pipeline using well-established methods, including: Trimming of low-quality bases and removal of sequencing adaptors with Fastp [43]; Ribosomal RNA (rRNA) removal matching against the <u>SILVA rRNA database</u> [44]; Transcriptome assembly using a range of tools, including Trinity, SPAdes, Trans-ABySS and SOAPdenovo-Trans, with a range of k-mers [45, 46]; Collapsing redundant transcripts with <u>EvidentialGene</u>; Annotation of the assemblies against UniProt databases; Building a <u>Trinotate</u> database for each assembly and generating an annotation. Since TransPi is not actively maintained, several issues were encountered in the installation and running of the pipeline in newer Nextflow versions and container systems (Apptainer). A modified version of TransPi was therefore used (available at https://github.com/IdoBar/TransPi), refer to https://idobar.github.io/QX_bioinfo_analysis/ for detailed description of the bioinformatics analysis, including specific details and code snippets of the modifications made to the pipeline.

Assembly completeness was determined using BUSCO [47] using the Metazoa database (Metazoa_Odb10) to assess the presence of highly-conserved single copy orthologs. The quality and coverage of the transcriptomic data was further assessed using TransRate [48].

#### 2.5.2 Transcriptome Annotation

The transcriptome assembly was annotated initially by TransPi, using homology searches (BLAST [49] and DIAMOND, [50]) against the Uniprot_Sprot protein database, as well as using HMM search against the Pfam database of protein families. In addition, signal peptides, ribosomal genes and transmembrane topology were predicted with SignalP v4.1[51], Rnammer and TMHMM [52], respectively [53].

#### 2.5.3 Gene Ontology and Functional Annotation

In addition to the annotations performed with TransPi, the assembled transcripts were annotated against the non-redundant nucleotide database of the NCBI database (nt) using BLASTn (v2.16.0) homology search to achieve more accurate species-specific annotations. To reduce computation time, we used <u>nf-blast</u>, a custom Nextflow workflow that splits the input transcriptome and processes them in parallel on the HPC cluster. The same tool was used to annotate the predicted proteins derived from the transcriptome using DIAMOND (v2.1.9) against the NCBI non-redundant protein database (nr). The predicted proteins were functionally annotated with InterProScan v5.66-98.0 [54] to assign protein families, motifs and ontologies to assist with transcript-to-gene annotation. Gene Ontology (GO) annotation terms were extracted from the InterProScan result tables.

#### 2.5.4 Differential Gene Expression Analysis

Data processing for transcript count generation were performed on Galaxy Australia [55]. The trimmed and corrected RNA-Seq reads that were used to assemble the transcriptome were then aligned back to the transcriptome using BWA-MEM2 (v2.2.1+galaxy1[56]) using default parameters. Transcript counts were generated from the alignments by featureCounts (v2.0.8+galaxy0[57]). Quality-trimming and alignment statistics were aggregated into a single report with MultiQC (v1.27+galaxy0[58]). The count tables were imported into R and DESeq2 (v1.42.1[59]) was used for differential gene expression analysis. Count data was imported into a DESeqDataSet object, and normalisation was performed using the default method in DESeq2. The design matrix included infection status as a factor, and differentially expressed genes were identified using the Wald test. The resulting p-value was adjusted to account for multiple testing using the Benjamin-Hochberg algorithm [60] to control the false discovery rate (FDR). Genes with an absolute log2 fold change (|log_2_FC|) > 2 and an adjusted p-value ≤ 0.05 were considered as significantly differentially expressed.

GO term enrichment analysis was performed using the clusterProfiler R package (v4.10.1[61]) on the identified differentially expressed genes. However, due to the poor protein annotation rates for molluscs (17%, see Results section 3.1), no GO terms were found to be enriched and therefore the GO terms were summarised to identify the most frequent terms in the biological process (BP) and molecular function (MF) categories. Lollipop plots visualising the most frequent annotations in all differentially expressed genes were generated using ggplot2.

#### 2.5.5 Data processing and visualisation

RNA-Seq data analysis and transcriptome assembly were performed on ‘Gowonda’, the Griffith University High Performance Computing cluster and ‘Bunya’, the University of Queensland High Performance Computing cluster. Downstream analyses were performed in the R environment for statistical computing (v4.1) [35, 36]. Various R packages from the tidyverse (v2.0.0) [37] along with the janitor package (v2.2.0) [38] were used for data cleaning, structuring, summarising and visualising. All code utilised for the analysis is available on Zenodo at https://doi.org/10.5281/zenodo.14564721 [39] (see <u>Data Availability</u> section).

A Principal Component Analysis (PCA) plot was generated to assess sample clustering based on normalised expression data. A heatmap of the top 500 differentially expressed genes was created using the pheatmap package (v1.0.12[62]), with hierarchical clustering applied to both genes and samples. A volcano plot depicting the relationship between fold change and statistical significance was generated using the EnhancedVolcano package (v1.13.2[63]).

## 3. Results and Discussion

### 3.1 Confirmation of *M. sydneyi* infection in samples

The presence and absence of *M. sydneyi* infection in *S. glomerata* samples was confirmed via PCR and histological tissue smears. PCR results were assessed based on amplicon brightness (Fig. 2A), as a previous correlation was observed between band intensity and the number of *M. sydneyi* cells in infected tissue (Fig.2B).

**Fig. 2.**
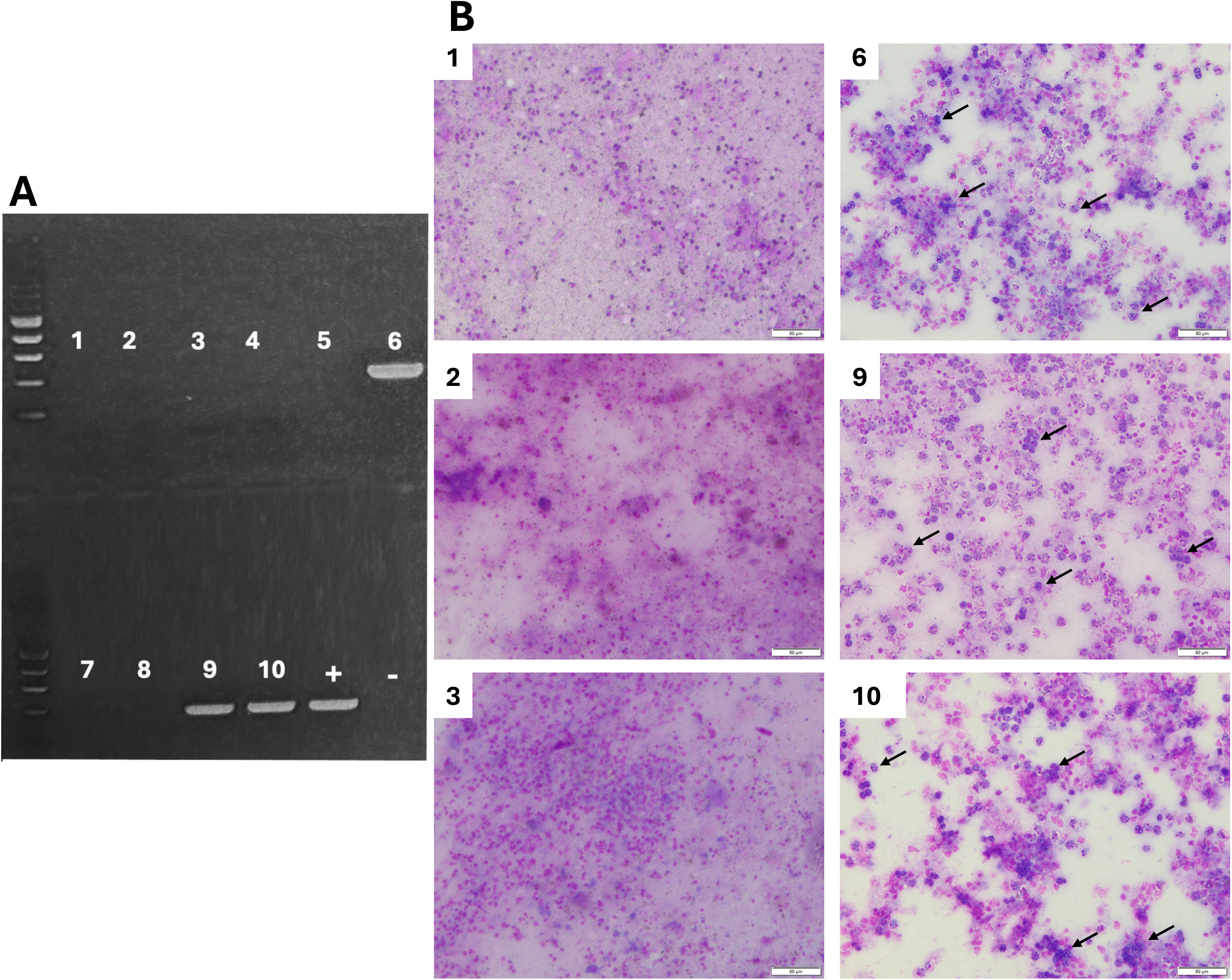
*M. sydneyi* infection diagnosis. (A) Gel electrophoresis of PCR amplicons generated using the Leg1 and Pro2 primers on *S. glomerata* digestive gland DNA. A 100bp ladder was used to aid visualization. Lanes 1-3: uninfected *S. glomerata* individuals 1, 2, and 3, respectively. Lanes 6, 9 and 10: *M. sydneyi* infected *S. glomerata* individuals 1, 2, and 3, respectively. Lanes 4,5,7,8: uninfected *S. glomerata* individuals that were not included for sequencing. +: positive control of *S. glomerata* confirmed to have *M. sydneyi* infection via PCR, histology and sequencing. -: no template control. (B) *S. glomerata* digestive gland histology tissue smears on individuals chosen for transcriptomic sequencing, scale bars = 50µm. 1-3: uninfected *S. glomerata* individuals 1, 2, and 3, respectively. 4-6: *M. sydneyi* infected *S. glomerata* individuals 1, 2, and 3, respectively. Black arrows indicate presporulating sporangiosori.

### 3.2 Transcriptome assembly and differential gene expression

RNA sequencing generated 35-39 million paired-end reads per sample, with read duplication rate ranging between 25% - 38%. On average, 94% of the reads per sample (equivalent to 9.9 million bases) had a phred score of over Q30 (Table 1). The assembled transcriptome of the *S. glomerata* digestive gland produced a total of 216,914 transcripts, with an N50 value of 1,235 bp, indicating the median contig length of the assembly’s longest 50% of the bases. The N10 value, representing the length above which 10% of the total assembled bases are found, was 3,713 bp, highlighting the presence of longer, well-assembled contigs. The assembly had a median contig length of 460 bp, and the overall GC content of the transcriptome was 36.8%, reflecting the nucleotide composition.

**Table 1.** General statistics from sequencing results and the alignment of raw reads against the *S. glomerata* transcriptome generated in this study.

| Sample Name | Condition | % GC<sup>a</sup> | % Read alignment to reference<sup>b</sup> | % Dups<sup>c</sup> | >Q30 bases (x10<sup>9</sup>)<sup>d</sup> | Million paired end reads |
| --- | --- | --- | --- | --- | --- | --- |
| NN01-561 | Uninfected 1 | 43.50% | 98.80% | 29.20% | 10.578 | 39.2 |
| NN01-562 | Uninfected 2 | 43.50% | 98.90% | 38.40% | 9.9 | 36.6 |
| NN01-563 | Uninfected 3 | 43.50% | 98.90% | 30.00% | 9.862 | 37 |
| NN01-567 | Infected 1 | 39.30% | 59.40% | 25.40% | 9.599 | 35.5 |
| NN01-579 | Infected 2 | 39.20% | 50.50% | 27.90% | 9.755 | 36.6 |
| NN01-580 | Infected 3 | 41.40% | 82.40% | 32.70% | 9.989 | 37.3 |

**a** - % GC refers to the percentage of guanine (G) and cytosine (C) bases in the sequenced RNA sequences. **b** - % Dups refers to the percentage of duplicate reads per sample. **c** – Percentage of reads aligned to the *S. glomerata* de-novo assembled transcriptome. **d** – number of bases (billions) over Q30 after filtering.

The BUSCO analysis of the de novo transcriptome assembly for *S. glomerata* digestive gland indicated a high level of completeness, with 95.3% of the expected core genes identified (Table 2). A small fraction (1.9%) of the genes were found to be fragmented, and 4.7% were missing from the assembly, indicating minimal loss of expected gene content. 74.3% of the BUSCO genes were duplicated, suggesting the transcriptome assembly is likely to contain some level of redundancy and duplication due to assembly and sequencing inaccuracies. Overall, the results indicate that the transcriptome assembly is of high quality, with good representation of the gene content.

**Table 1.** BUSCO completeness scores for the de novo *S. glomerata* transcriptome assembly.

| DESCRIPTION | RESULTS |
| --- | --- |
| Complete and single-copy BUSCOs | 19.1% |
| Complete and duplicated BUSCOs | 74.3% |
| Fragmented BUSCO genes | 1.9% |
| Missing | 4.7% |

**Table 2:** Summary of differentially expressed genes between infected and uninfected *S. glomerata* samples

| DESCRIPTION | TRANSCRIPTS |
| --- | --- |
| Number of DEG | 2,200 |
| Number of annotated DEG | 380 |
| Number of DEG upregulated in infected tissue | 1,894 |
| Number of DEG upregulated in healthy tissue | 306 |

Alignment of the reads to the assembled transcriptome revealed that the uninfected samples had a high percentage of reads that aligned to the reference *S. glomerata* digestive transcriptome (approximately 98.9%), while infected samples had much lower alignment rates, indicating the presence of parasite-derived reads. Infected samples 1 and 2 had 59.4% and 50.5% of reads align to the reference genome, whereas infected sample 3 showed a higher alignment rate of 82.4%, falling between the uninfected and other infected samples. Although all three samples appeared to have similar levels of infection based on PCR results and histology tissue smears (Fig. 2), the distribution of *M. sydneyi* cells throughout infected digestive gland tissue could vary. Therefore, the tissue sampled from infected sample 3 for RNA extraction may have had a lower parasite load than initially thought. Assessment of GC content in the different samples suggests this may be the case (Supplementary Fig. 2).

### 3.3 Differential gene expression

Of the total 216,914 transcripts analysed, 2,200 were found to be differentially expressed between infected and uninfected samples, emphasizing the substantial and wide-ranging molecular responses to infection (Table 3 and Fig. 3C). There were clear patterns of expression between infected and uninfected *S. glomerata* individuals, demonstrating the substantial impact *M. sydneyi* infection has on *S. glomerata* gene expression profiles in the digestive tissues (Fig. 3A). This distinction was further supported by the PCA plot, where the samples from the two experimental groups clustered with a clear separation between infected and uninfected samples (Fig. 3B).

**Fig. 3.**
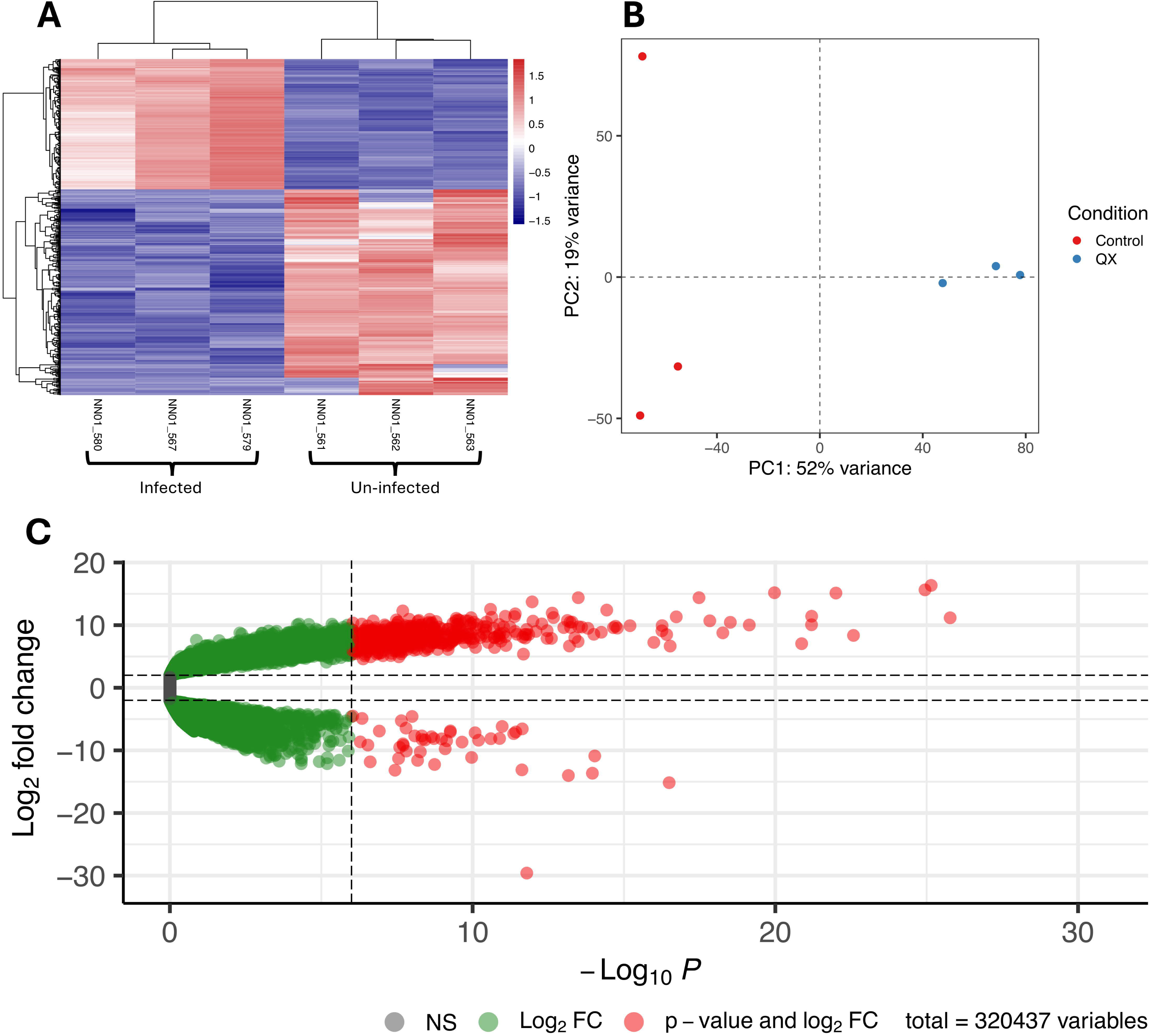
Differential gene expression in infected and uninfected *S. glomerata*. (A) Heatmap of the top 500 differentially expressed genes in *M. sydneyi* infected *S. glomerata* digestive tissue. (B) PCA plot of infected and un-infected samples. (C) Volcano plot highlighting the log2fold change and p-value of differentially expressed genes. NS (non-significant) indicates genes that had an absolute log_2_FC less than 2 in differential gene expression analysis. Transcripts indicated in green had an absolute log_2_FC greater than 2, and transcripts indicated in red had both an absolute log_2_FC greater than 2 and an adjusted p-value less than 0.05 (considered significant in our study).

Previous studies have shown that transcriptional responses to pathogens can be large, and that they are dependent on the stage of infection. For example, a study on *M. gigas* infected with *Vibrio alginolyticus* identified 1,543 DEGs at 6 hours post-infection, increasing to 2,483 DEGs at 48 hours [64], demonstrating that our findings align with prior observations and are within a comparable range. The high number of DEGs found in our study could be attributed to the severity and progress of *M. sydneyi* infection in the oysters, with substantial tissue damage and animals possibly close to mortality. Pathogen-induced stress and tissue damage during later stages of infection likely amplify DEGs as the host attempts to counteract infection and repair cellular damage.

### 3.4 Gene ontology classifications

Only 17% of the differentially expressed transcripts were successfully annotated (Table 3). Annotation rates in non-model organisms such as molluscs, including *S. glomerata*, are often limited due to the scarcity of genomic resources, highlighting the need for further functional characterisation efforts in this taxonomic group [65, 66].

The DEGs were analysed by GO term classification to explore whether certain biological processes or molecular functions are associated with *M. sydneyi* infection. GO biological processes that included more than four differentially expressed genes were visualised and are shown in Fig. 4A. The GO biological process analysis revealed a variety of responses associated with infection in oysters. These included genes involved in the activation of inflammatory responses, indicating the presence of a continued immune reaction despite the late stage of infection. Xenobiotic metabolic processes were observed to have four DEG within the group, suggesting that detoxification processes may play a vital role as a defence mechanism against infection-induced stress (Fig. 4A). Additionally, the presence of genes related to serine-type endopeptidase activity (5 DEG) and oxidoreductase activity (16 DEG) suggests involvement in proteolysis and redox processes (Fig. 4B). This suggests that QX infection induces protein degradation, turnover, and alterations in redox balance, which are a likely part of the host’s response to cellular stress. These changes may represent a defensive mechanism or processes involved in immune regulation. Upregulation of genes linked to cell matrix adhesion and nuclear envelope organisation in infected tissues likely reflects the poor condition of the infected digestive tissues (Fig. 2B, 3). This indicates that the infection impacts cellular integrity and signalling, disrupting normal cellular functions and structure [67, 68].

**Fig. 4.**
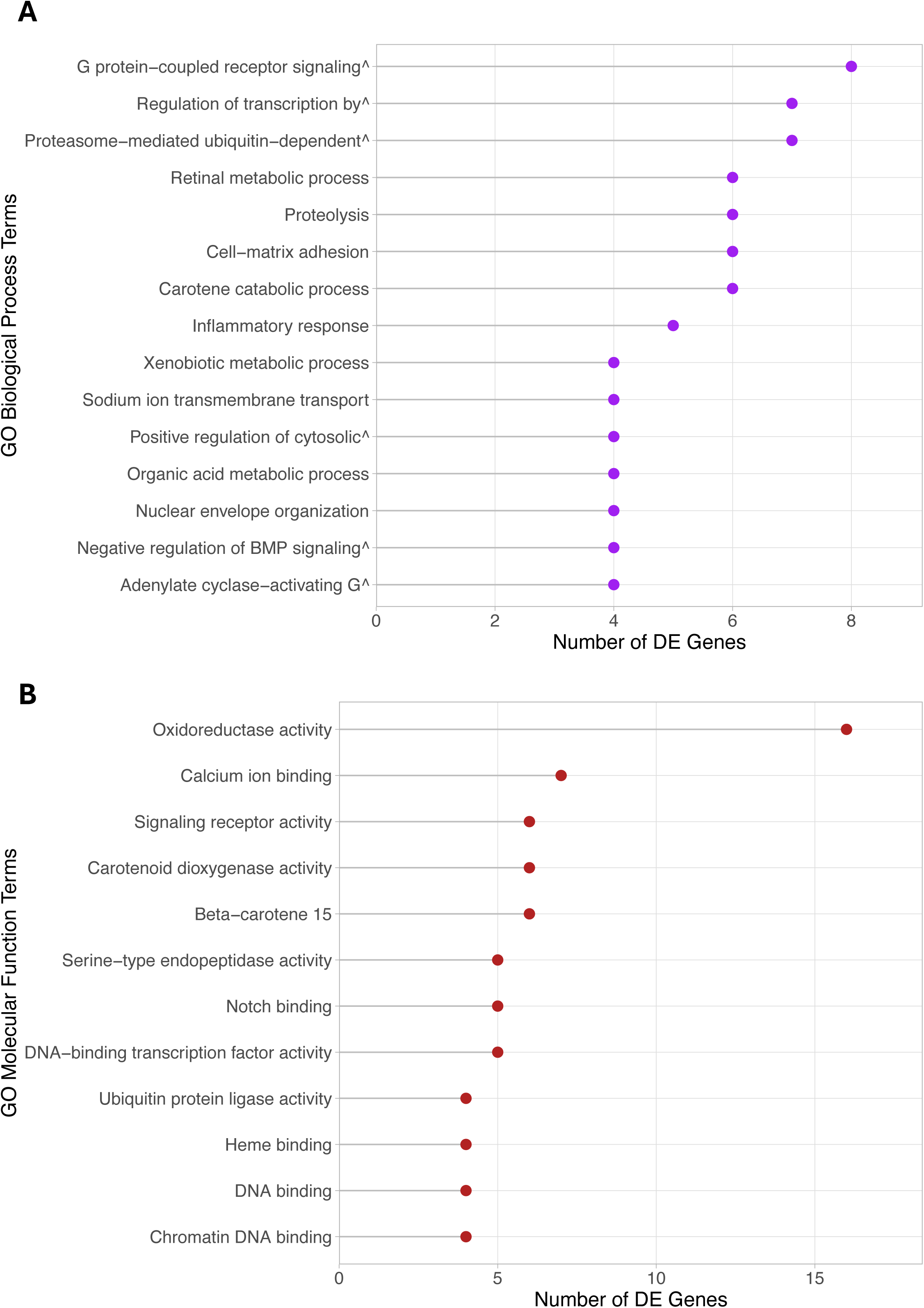
Gene ontology annotations of genes that are differentially expressed in response to *M. sydneyi* infection in *S. glomerata*. (A) GO biological process terms associated with four or more DEGs. (B) GO molecular function terms associated with four or more DEGS.

A similar approach was applied to GO molecular functions, focusing on categories containing a minimum of four differentially expressed genes. Fig. 4B highlights the involvement of transcription and chromatin-related activities, including terms such as DNA-binding transcription factor activity, chromatin binding, and DNA binding, supporting our observation that the infection causes widespread changes in gene regulation [69, 70].

### 3.5 Functional Annotation of Differentially Expressed Genes

Despite the low annotation rates, some interesting differentially expressed gene were identified that appear relevant to *S. glomerata’s* response to *M. sydneyi* infection. The annotated genes with the largest transcriptional changes in *S. glomerata* infected with *M. sydneyi* are shown in Fig. 5 and include galectin, cytochrome P450, ecSOD, fibrinogen-related proteins, and G-coupled receptors. The magnitude of differential gene expression likely reflects the complex interaction between host immune defences, potential evasion strategies employed by the parasite, and the significant tissue damage observed during infection. Host immune defences may activate specific pathways to target and neutralize the parasite, while the parasite simultaneously employ evasion mechanisms, such as suppressing immune signalling, altering host cell processes, or avoiding recognition altogether. Several of these highly differentially expressed genes have also been identified in previous studies focusing on transcriptomic responses in bivalves to parasitic infections. Among these genes, notable examples include galectins, cytochrome P450 family genes and extracellular superoxide dismutase (ecSOD). These genes represent diverse functional categories. The identification of the same response genes in different bivalves infected with different parasites suggests that the host response pathway may be highly conserved across different taxa.

**Fig. 5.**
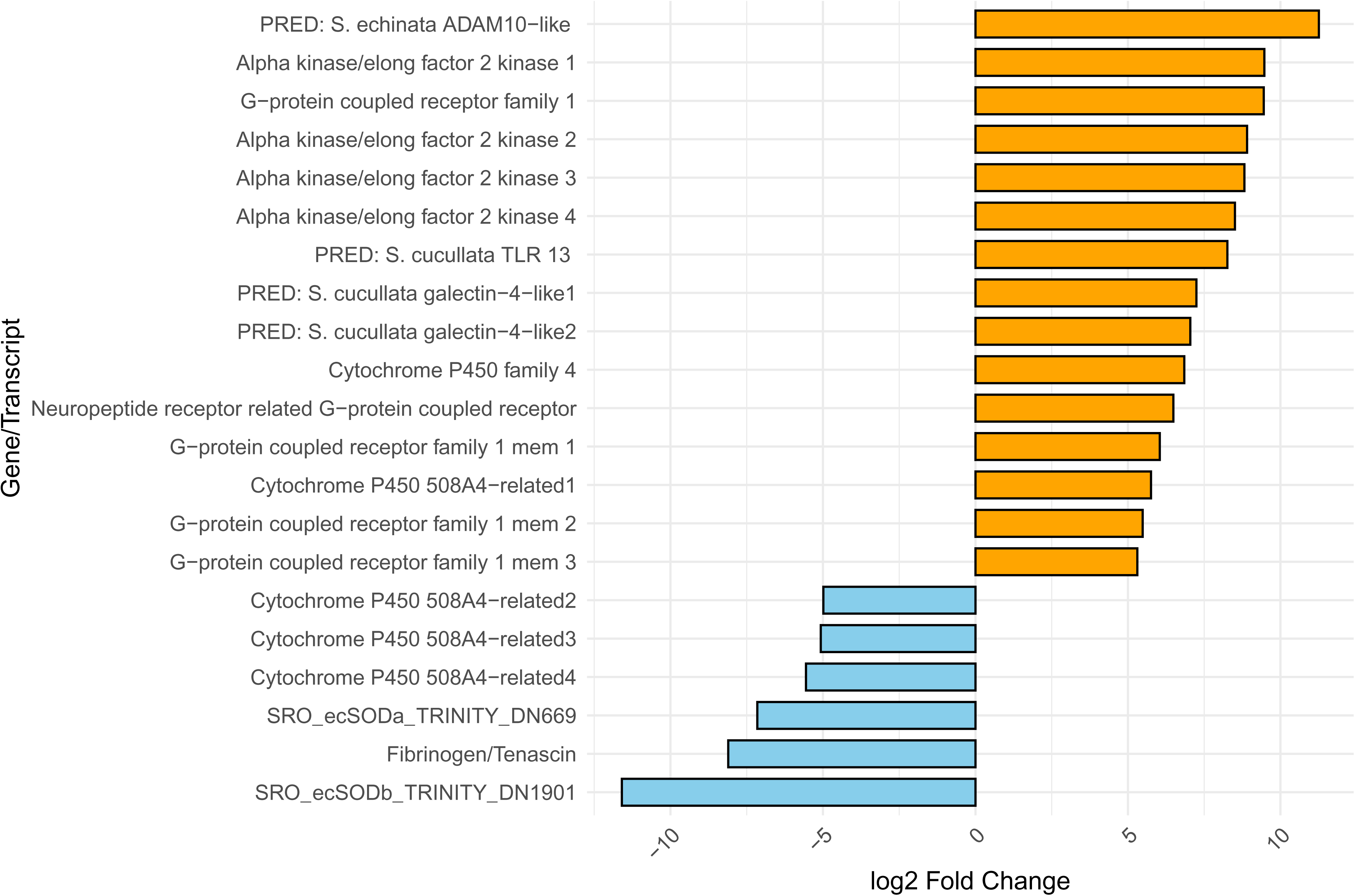
Annotated differentially expressed genes that had a minimum log_2_fold change of 5 or -5 between infected and uninfected *S. glomerata* digestive gland tissue. Yellow bars indicate up-regulated genes and blue bars indicate down-regulated genes in *S.glomerata* infected with *M. sydneyi*.

#### 3.5.1 Galectin

Galectins are a family of lectins distinguished by their specific binding of β- galactosides and their conserved sequence motif within the carbohydrate recognition domain [72]. In *S. glomerata* several predicted galectins are upregulated during *M. sydneyi* infection (Fig. 5). Similar findings in other bivalve species (including *O. edulis, M. gigas,* and *Ruditapes philippinarum*) have indicated that galectins are involved in pathogen recognition, regulation of immune cell activity, and binding to specific surface glycans on micro-organisms that facilitates immune responses [11, 31, 73]. For example, galectins identified in *C. virginica* have been shown to target and bind to external glycans on bacteria and parasites [74, 75]. This suggests that the upregulation of galectin-4-like in *S. glomerata* during *M. sydneyi* infection may enhance the oyster’s ability to identify and respond to the parasite by promoting surface glycan binding of the parasite to host immune cells, potentially regulating downstream immune signalling pathways.

#### 3.5.2 Cytochrome P450

Cytochrome P450 (CYP450) represents a diverse superfamily of hemoproteins found in all biological kingdoms, characterised by a heme cofactor that facilitates a wide range of catalytic functions [76–78]. CYP450 enzymes are involved in a range of physical processes such as the metabolism of endogenous and xenobiotic compounds, the breakdown of environmental toxins, and hormone biosynthesis [76–78]. They mediate various reactions, including hydroxylation, peroxidation, epoxidation and reduction, contributing to detoxification and cellular protection mechanisms [77].

In some bivalves, CYP450 family members are known for their involvement in cellular protection mechanisms and post-phagocytosis degradation, most notably oxidative metabolism [79]. The presence of multiple CYP450 family members within both upregulated and downregulated DEGs suggests a complex regulatory response to *M. sydneyi* infection. There was a significant up-regulation of some CYP450 family members, with a logfold change of approximately 6.85 (Fig. 5), congruent with the activation of enhanced metabolic or detoxification activities as a possible response to infection [80]. This expression pattern aligns with results seen in other bivalve studies investigating host-pathogen interactions, in which CYP450 superfamily genes were linked to oxidative metabolism and detoxification during infection. For example, in *Mercenaria mercenaria* CYP450 was significantly upregulated following Quahog parasite unknown (QPX) infection, where it was proposed that the gene plays a role in detoxifying damaging compounds produced during the initial immune response [81]. Upregulation of CYP450 was associated with initial stages of *B. ostreae* infection in *O. edulis*, potentially also as a response to oxidative stress [79]. However, our study revealed that three of the CYP450 family member genes were observed to be downregulated, indicating possible suppression of the metabolic and detoxification processes they are involved in. This could be a means to conserve energy or avoid excessive oxidative damage. The varying expression levels of CYP450 genes suggests they have multifaceted roles in *S. glomerata’s* defence against *M. sydneyi*. This likely reflects the host’s approach to balancing immune defence, metabolic adaption and oxidative stress.

#### 3.5.3 Extracellular superoxide dismutase

In bivalves, ROS is utilised as a key defence mechanism during phagocytosis. ecSOD is a key enzyme involved in this process, converting superoxide anions to hydrogen peroxide, an anti-parasitic compound [82, 83]. Previous research has shown elevated expression of ecSOD genes in disease-resistant *S. glomerata* and *O. edulis* [31, 82]. It was also recently shown that *S. glomerata* possesses two copies of the ecSOD gene, ecSODa and ecSODb (with ecSODa corresponding to the previously investigated *S. glomerata* ecSOD; the role of ecSODb is unknown) [84]. Prior studies on ecSODa expression in *S. glomerata* revealed the gene to be non-inducible upon bacterial infection with *Vibrio anguillarum*, indicating that expression of ecSOD is not modulated in response to infection [30, 85].

In this study, we observed a significant downregulation of all isoforms of both ecSOD genes in *S. glomerata* infected with *M. sydneyi* (Fig. 5, 6). This could imply a weakened oxidative stress response to infection, potentially compromising the host’s ability to manage ROS, or it may reflect the host’s efforts to moderate immune responses by reducing ROS production, potentially minimizing self-inflicted oxidative damage to tissues and cells. The moderation of ROS production may serve to preserve energy and resources during prolonged immune responses, allowing the host to allocate resources more efficiently to other defence mechanisms [86, 87]. Conversely, it could indicate the parasite’s successful suppression of host defences, allowing it to evade ROS-mediated destruction [79]. Parasite suppression of host responses has been proposed in *B. ostreae* infection in *M. gigas* and *O. edulis,* where a downregulation of ecSOD during infection was associated with reduced phagocytic activity, potentially increasing the parasite’s chances of survival [11, 18, 79]. Similarly, *M. sydneyi* infection in *S. glomerata* has been linked to the suppression of the phenoloxidase cascade, a critical immune pathway involved in pathogen encapsulation and melanisation [19, 24, 88]. This suppression is also associated with reduced phagocytic activity, further facilitating parasite persistence within the host [19, 24]. These findings suggest that immune suppression mechanisms employed by the parasite may play a role in disease progression and host susceptibility.

**Fig. 6.**
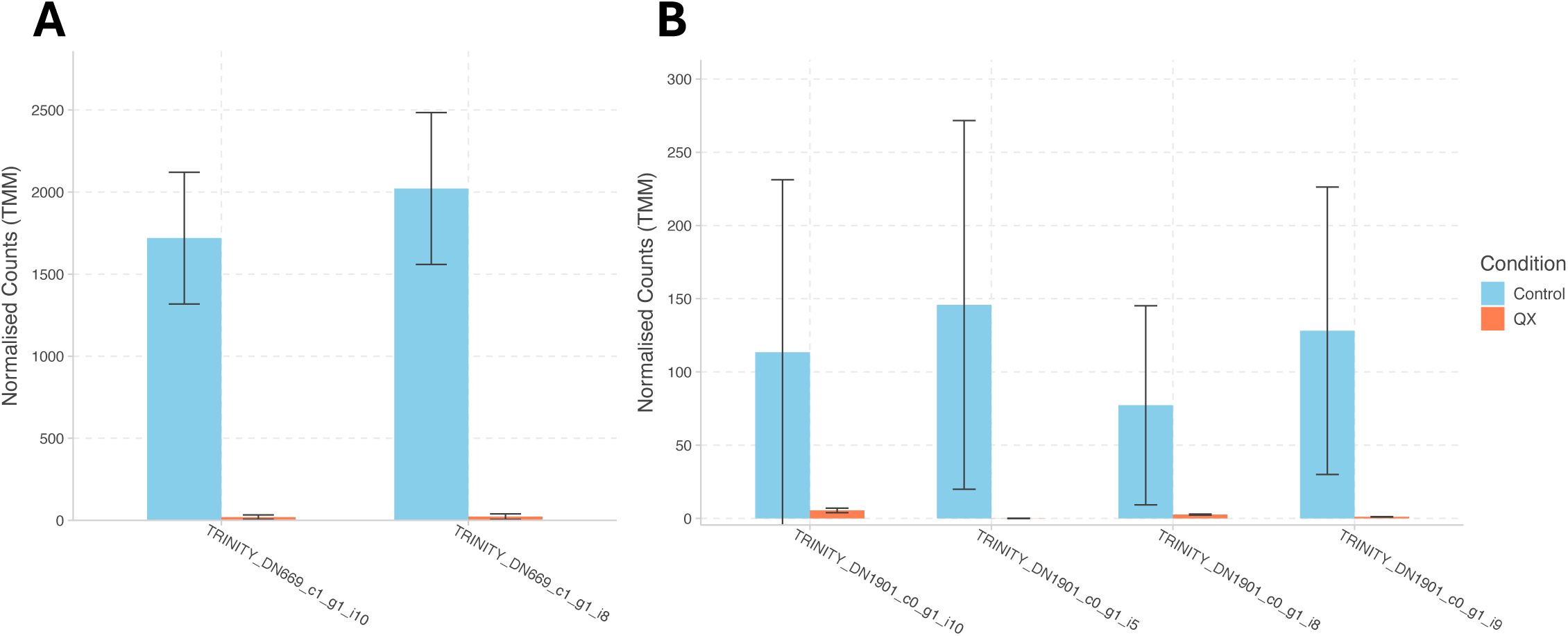
Gene expression of ecSOD transcripts in *M. sydneyi* infected (QX) and uninfected (control) *S. glomerata* tissue. There are multiple isoforms of ecSODa and ecSODb within the transcriptome assembly; all isoforms show downregulation in infected tissue. (A) Normalised counts of *S. glomerata* ecSODa transcripts. (B) Normalised counts of *S. glomerata* ecSODb transcripts.

Previous studies observed a downregulation of Prx6 in selectively bred disease resistant oysters, alongside an upregulation of ecSOD, suggesting a coordinated oxidative stress response [82]. Our findings suggest that Prx6 expression may be reduced in *S. glomerata* infected with *M. sydneyi*, as indicated by an apparent decrease in normalised Prx6 counts in infected samples compared to uninfected samples (Fig. 7). However, this decrease was not statistically significant in DGE analyses, making it unclear whether Prx6 suppression plays a major role in the host response to infection. Given that ecSOD plays a key role in ROS detoxification, its suppression in infected oysters could impair the host’s ability to regulate oxidative stress, potentially making them more susceptible to tissue damage and pathogen persistence. If the expression of Prx6 is also reduced, this could further compromise the oxidative stress response, weakening host defences against *M. sydneyi* infection.

**Fig. 7.**
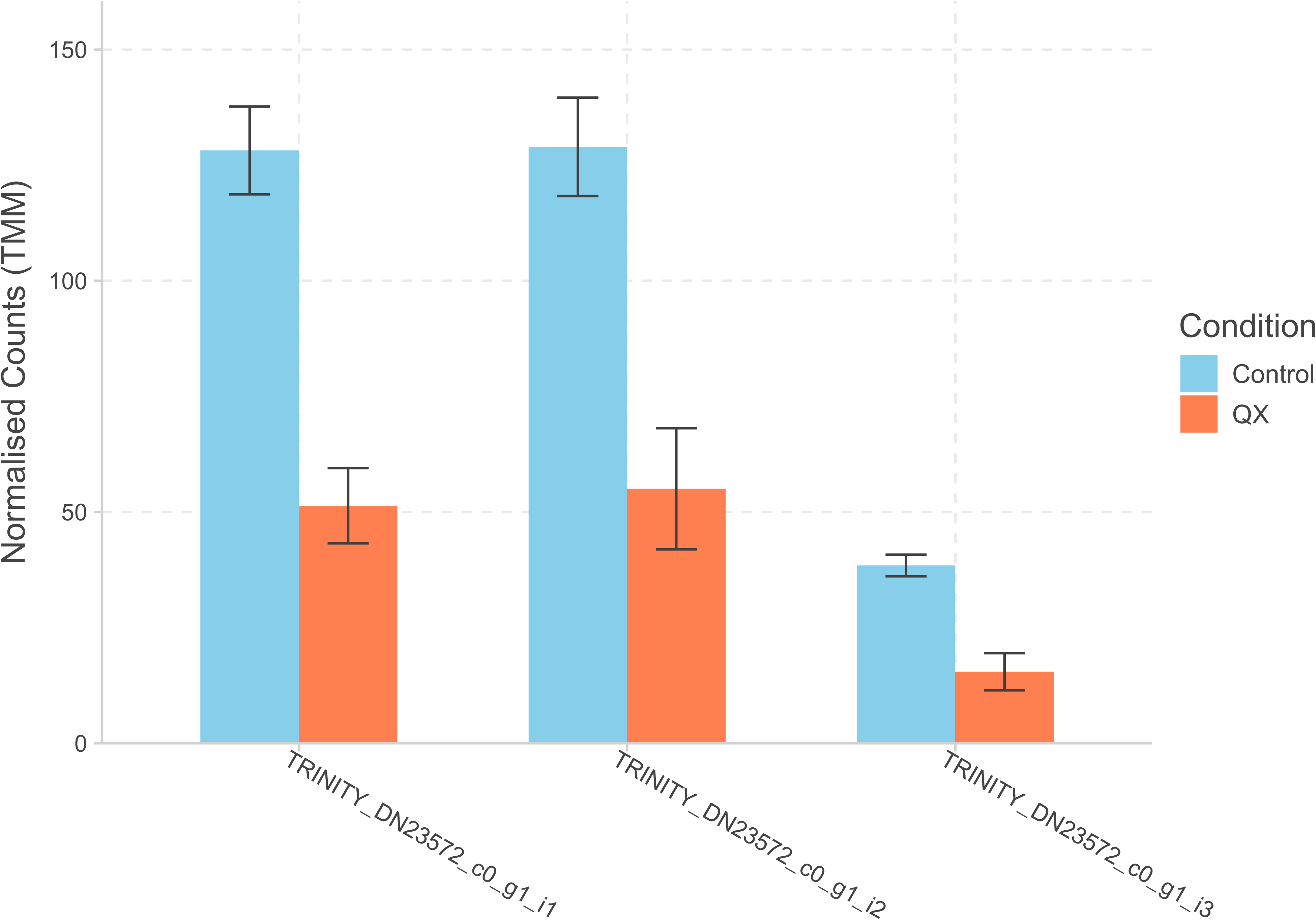
Gene expression of Prx6 transcripts in *M. sydneyi* infected (QX) and uninfected (control) *S. glomerata tissue*. There are three isoforms of Prx6 within the transcriptome assembly; all isoforms show downregulation in infected tissue.

#### 3.5.4 Fibrinogen-related protein family

Fibrinogen-related proteins (FREPs) are proteins that contain a fibrinogen-related domain (FReD). They are pattern recognition receptors with vital roles in invertebrate innate immune responses [89, 90]. The gene family is highly diverse and encodes multiple protein types that are typically upregulated following immune stimulation [91, 92]. FREPs are able to bind to pathogen cells and precipitate parasite antigens [92]. Upregulation of FREPs in QPX resistant *M. mercenaria* clams led to the gene family being recognised as a potential marker for resistance due to its ability to increase defence responses against pathogens [71]. In contrast with previous findings, Fibrinogen/Tenascin was downregulated in *S. glomerata* infected with *M. sydneyi* (Fig. 5), suggesting a potential disruption in immune processes. This suppression could indicate the parasite’s interference with the host’s immune pathways, reducing the host’s ability to effectively detect and respond to the parasite. The downregulation of this gene in the present study could suggest an evasion strategy by *M. sydneyi* to undermine the hosts immune response.

#### 3.5.5 G-protein coupled receptors

G-protein coupled receptors (GPCRs) are essential for a multitude of immune processes, including the activation of transcription factors that regulate downstream inflammatory responses, such as NF-κB, CREB, and STAT3 [93, 94]. Several GPCRs were upregulated in *S. glomerata* tissue infected with *M. sydneyi* (Fig. 5). The most significant upregulation is seen in GPCR family 1 member with a log2fold change of 9.45. Other GPCR genes also display significant upregulation, with log2fold change values ranging from 5.31 to 6.04. The significant upregulation of multiple GPCRs in the host indicates that GPCRs play an active role in mediating the host’s immune response signalling. The upregulation seen could suggest an enhanced immune surveillance mechanism in oysters, as GPCRs assist in the migration and activation of immune cells at sites of infection [18]. Recent studies focusing on host-pathogen interactions between *M. gigas* and ostreid herpes virus, and *M. mercenaria* infection with QPX, found that GPCRs were associated with defence responses [83, 95]. Upregulation of these receptors may reflect the oyster’s attempt at boosting immune signalling pathways and managing infection-induced stress. While the upregulation of GPCRs suggests an attempt by the host to enhance immune surveillance and immune cell activation, the parasite may counteract these efforts by suppressing downstream signalling pathways, leading to an overall weakened immune response. Studies investigating host/pathogen interactions between *B. ostreae* and *M. gigas* revealed downregulation of GPCRs in lightly infected oysters, with further diminishment with heavier infection [96, 97]. This pattern highlights the parasite’s active suppression of these pathways as a strategy to evade host immune responses. Overall, the results underline the potential role of GPCRs in regulating oyster defence mechanisms and the importance of understanding host-pathogen molecular responses.

#### 3.5.6 The role of apoptosis in the *S. glomerata*- *M. sydneyi* interaction

Interestingly, apoptosis or necrosis-related genes (such as p53, Bcl-2 proteins, or caspases), were not found to be differentially expressed in our analysis. The stable expression of these genes suggests that programmed cell death may not be a prominent immune response at this stage of the infection or a significant feature in the host’s immune response to *M. sydneyi* infection. Nevertheless, studies indicate that immune challenges can induce apoptosis through both pre-existing pathways (caspase activation) and changes in gene expression, particularly under prolonged or complex stress conditions [98, 99]. As mentioned, previous research on *C. virginica* infected with *P. marinus* has demonstrated the parasite’s ability to survive within host hemocytes by modulated apoptotic pathways, ultimately facilitating infection persistence [8, 100]. Specifically, *P. marinus* has been shown to upregulate antioxidant enzymes such as superoxide dismutase (SOD) and Prx6, which help suppress the production of ROS and delay apoptosis in host cells [101, 102]. This strategy allows the parasite to evade host defences and continue intracellular replication.

Apoptotic responses in oyster hemocytes exposed to *P. marinus* have been observed to follow a dynamic pattern, with an early increase in apoptosis followed by suppression over time [8, 100]. This temporal regulation indicates that the parasite may actively manipulate apoptotic pathways to establish infection. The intrinsic mitochondrial apoptosis pathway, which involved key regulators such as Bcl-2, plays a significant role in this process by potentially inhibiting cytochrome-c release and delaying cell death [103, 104]. Studies have further revealed that anti-apoptotic genes are upregulated in virulent cultures of *P. marinus,* suggesting that the parasite utilises these mechanisms to suppress apoptosis and enhance survival in the host [105, 106].

In contrast, our findings imply that *M. sydneyi* may employ a different strategy to evade host immune responses, potentially bypassing the need to regulate apoptosis at the transcriptional level. The absence of significant changes in apoptosis-related gene expression suggests that apoptosis may not be a critical component of the immune response to *M. sydneyi*, or that the response occurs through non-transcriptional modifications. Further investigation, including protein-level analysis and temporal studies are needed to better understand whether apoptosis plays a role at earlier or later stages of infection, or in specific tissues, as our comparisons were limited to a specific time point.

#### 3.5.7 Conserved molecular responses to parasite infection in bivalves

The findings of this study highlight potential shared defence strategies across bivalve species, suggesting that fundamental immune responses may be conserved across a range of pathogens rather than being specific to ascetosporean parasites. Comparing the results of this study with that of *O. edulis* infected with *B. ostreae* reveals the possibility of shared defence strategies between the *O. edulis* and *S. glomerata*, including the central role of glycan-binding proteins such as FREPs and lectins in pathogen recognition [18]. Glycan-binding proteins have been reported in bivalve immune systems as key components of non-self-recognition, mediating interactions with pathogen glycans and activating downstream immune pathways. In *M. gigas,* lectins were observed to bind to PAMPs and play a significant role in immune surveillance against multiple pathogens, underscoring their conserved function across bivalve species [12].

Oxidative stress regulation also appears to be a conserved immune mechanism across bivalves, with a downregulation of ecSOD observed in both *O. edulis* and *S. glomerata* [79]. This down regulation may reflect a parasite-driven suppression of oxidative stress pathways to evade host immune responses, or a host-mediated response to limit further tissue damage from excessive ROS. Previous investigations on *M. galloprovincialis* during bacterial infections emphasized the dual role of oxidative stress pathways in balancing pathogen elimination and host tissue preservation. The increased activity of antioxidant enzymes, such as catalase and GST, reflects an active response by the digestive gland to manage the heightened production of ROS triggered by bacterial challenges [107]. Specifically, infections with *Vibrio splendidus* and/or *Vibrio anguillarum* have been shown to enhance the activities of these enzymes, highlighting their necessity in mitigating oxidative damage and preserving cellular integrity whilst supporting immune defence responses [107]. The similarities in oxidative stress regulation across different bivalve-pathogen interactions suggest that these mechanisms represent a fundamental aspect of molluscan immunity.

Species-specific differences in immune responses were evident in the structural responses and signalling pathways observed between *M. gigas* and *S. glomerata*, suggesting potential adaptations to distinct pathogenic or environmental pressures, Previous studies noted that structural responses involving extracellular matrix remodelling were notable in *M. gigas,* suggesting an essential role in maintaining tissue integrity and facilitating immune cell migration during infection [108]. In contrast, *S. glomerata* demonstrated significant upregulation of GPCR pathways, which are known to mediate immune signalling.

## 4. Conclusions

This study highlights the significant molecular changes that *S. glomerata* undergoes when infected with *M. sydneyi*, including notable alterations in cellular defence mechanisms, oxidative stress management, and immune response genes. The upregulation of genes such as galectin-4-like and multiple GPCRs suggests their involvement in immune signalling and pathogen recognition, potentially reflecting the host’s attempt to increase or initiate immune surveillance strategies. Conversely, the significant changes in the expression of CYP450 family genes, ecSOD, FREPs and GPCRs imply that *M. sydneyi* may employ molecular strategies to suppress the host’s oxidative burst defences, thereby weakening the oyster’s ability to combat infection. These findings not only broaden our understanding of *S. glomerata’s* immune responses but also identify key genes that could serve as biomarkers for breeding disease-resistant oysters and provide valuable targets for investigating the molecular basis of oyster immunity in future studies. Comparative insights with other bivalves not only enhance our understanding of shared immune strategies but also reveal species-specific adaptation that could inform aquaculture practices and disease management. Future research should further explore the host-pathogen interactions and molecular mechanisms underpinning *M. sydneyi* infections, as these could reveal that strategies the parasite utilises to manipulate host immune responses for successful infection.

## List of abbreviations

QX: Queensland Unknown
PAMPs: pathogen associate molecular patterns
ROS: reactive oxygen species
MSX: multinucleated sphere unknown
EcSOD: extracellular superoxide dismutase
Prx6: peroxiredoxin 6
IK: interferon-Inhibiting Cytokine Factor
SOD: superoxide dismutase
TIMP: tissue inhibitor of metalloproteinase
ECM: extracellular matrix
FREPs: fibrinogen-related proteins
C1qDCs: complement C1q domain-containing proteins
CRD: carbohydrate recognition domain
CYP450: Cytochrome P450
QPX: Quahog parasite unknown
GPCR: G-protein couple receptor

## Acknowledgements

We thank T Prowse and A Prowse from the Queensland Oyster Company for providing us with oysters from their lease. We thank T Prowse, A Prowse, M Richardson and M Stefanek for support with sample and data collection. We thank K Roper for the development of the altered standard RNA extraction using Trizol or Tri-reagent protocol utilised for our RNA extractions. Computing resources for the bioinformatic analyses were kindly provided by Griffith University’s Gowonda HPC cluster and the University of Queensland’s Bunya HPC cluster.

## Author contributions: CRediT

N. N.: Conceptualisation, formal analysis, investigation, methodology, visualisation, writing – original draft, writing – review and editing. I. B.: Conceptualisation, investigation, methodology, supervision, writing – review and editing. C. M.: Conceptualisation, investigation, methodology, supervision, writing – review and editing.

## Data availability

The data used in the study are available on NCBI under BioProjectID: PRJNA1173877. http://www.ncbi.nlm.nih.gov/bioproject/1173877. All bioinformatic analysis code is available on GitHub at https://idobar.github.io/QX_bioinfo_analysis/. All downstream analysis code is available on Zenodo at https://doi.org/10.5281/zenodo.14564721.

## Figure Legends

Supplementary Figure 1. Histology tissue smears of *S. glomerata* digestive gland tissue infected with *M. sydneyi*. Black arrows indicated presporulating sporangiosori of *M. sydneyi*. (A) *S. glomerata* digestive gland tissue frozen at -80°C then thawed prior to tissue smears, visualization at 1000x magnification. (B) *S. glomerata* digestive gland fresh tissue smears, taken immediately after dissection, visualized at 400x magnification.

Supplementary Figure 2. GC content per sequence sample (forward and reverse), generated by MultiQC. Green indicates *M. sydneyi* positive samples (infected 1 and 2), orange represents *M. sydneyi* infected sample 3, and red represents uninfected samples 1 and 2. The peak for infected sample 3 falls between infected and uninfected samples, suggesting that the sample may have had a lower parasite load than infected samples 1 and 2.

## References

1. van der Schatte Olivier A, Jones L, Vay LL, Christie M, Wilson J, Malham SK. A global review of the ecosystem services provided by bivalve aquaculture. Reviews in Aquaculture. 2020;12:3–25.

2. Shumway SandraE, Davis C, Downey R, Karney R, Kraeuter J, Parsons J, et al. In praise of sustainable economies and environments. World Aquaculture. 2003;34.

3. Lam K, Morton B. Morphological and mitochondrial-DNA analysis of the Indo-West Pacific rock oysters (Ostreidae: Saccostrea species). Journal of Molluscan Studies. 2006;72:235–45.

4. Botta R, Asche F, Borsum JS, Camp EV. A review of global oyster aquaculture production and consumption. Marine Policy. 2020;117:103952.

5. Okon EM, Birikorang HN, Munir MB, Kari ZA, Téllez-Isaías G, Khalifa NE, et al. A Global Analysis of Climate Change and the Impacts on Oyster Diseases. Sustainability. 2023;15:12775.

6. Hiltunen Thorén M, OnuC-Brännström I, Alfjorden A, Pecková H, Swords F, Hooper C, et al. Comparative genomics of Ascetosporea gives new insight into the evolutionary basis for animal parasitism in Rhizaria. BMC Biology. 2024;22:103.

7. Sm B, D H, Gr M. Infectivity of *Mikrocytos mackini*, the causative agent of Denman Island disease in Pacific oysters *Crassostrea gigas*, to various species of oysters. Diseases of Aquatic Organisms. 1997;29:111–6.

8. Hughes FM, Foster B, Grewal S, Sokolova IM. Apoptosis as a host defense mechanism in *Crassostrea virginica* and its modulation by *Perkinsus marinus*. Fish & Shellfish Immunology. 2010;29:247–57.

9. Burreson EM, Ford SE. A review of recent information on the Haplosporidia, with special reference to *Haplosporidium nelsoni* (MSX disease). Aquat Living Resour. 2004;17:499–517.

10. Laing I, Dunn P, Peeler EJ, Feist SW, Longshaw M. Epidemiology of Bonamia in the UK, 1982 to 2012. Diseases of Aquatic Organisms. 2014;110:101–11.

11. Cocci P, Roncarati A, Capriotti M, Mosconi G, Palermo FA. Transcriptional Alteration of Gene Biomarkers in Hemocytes of Wild *Ostrea edulis* with Molecular Evidence of Infections with Bonamia spp. and/or *Marteilia refringens* Parasites. Pathogens. 2020;9:323.

12. Guo X, He Y, Zhang L, Lelong C, Jouaux A. Immune and stress responses in oysters with insights on adaptation. Fish & Shellfish Immunology. 2015;46:107–19.

13. Söderhäll K, Cerenius L. Role of the prophenoloxidase-activating system in invertebrate immunity. Current Opinion in Immunology. 1998;10:23–8.

14. Roberts SB, Sunila I, Wikfors GH. Immune response and mechanical stress susceptibility in diseased oysters, *Crassostrea virginica*. J Comp Physiol B. 2012;182:41–8.

15. Wang L, Song X, Song L. The oyster immunity. Developmental & Comparative Immunology. 2018;80:99–118.

16. Akira S, Uematsu S, Takeuchi O. Pathogen recognition and innate immunity. Cell. 2006;124:783–801.

17. Morga B, Arzul I, Faury N, Renault T. Identification of genes from flat oyster Ostrea edulis as suitable housekeeping genes for quantitative real time PCR. Fish & Shellfish Immunology. 2010;29:937–45.

18. Ronza P, Cao A, Robledo D, Gómez-Tato A, Álvarez-Dios JA, Hasanuzzaman AFM, et al. Long-term affected flat oyster (*Ostrea edulis*) haemocytes show differential gene expression profiles from naïve oysters in response to *Bonamia Ostreae*. Genomics. 2018;110:390–8.

19. Butt D, Raftos D. Phenoloxidase-associated cellular defence in the Sydney rock oyster, *Saccostrea glomerata*, provides resistance against QX disease infections. Developmental & Comparative Immunology. 2008;32:299–306.

20. Aladaileh S, Rodney P, Nair SV, Raftos DA. Characterization of phenoloxidase activity in Sydney rock oysters (*Saccostrea glomerata*). Comparative Biochemistry and Physiology Part B: Biochemistry and Molecular Biology. 2007;148:470–80.

21. Nell J. The history of oyster farming in Australia. Marine Fisheries Review. 2001;83:14–22.

22. Steven A, Mobsby D, Curtotti R. Australian Fisheries and Aquaculture Statistics 2018. Canberra: ABARES; 2020.

23. Raftos DA, Kuchel R, Aladaileh S, Butt D. Infectious microbial diseases and host defense responses in Sydney rock oysters. Frontiers in Microbiology. 2014;5.

24. Peters R, Raftos DA. The role of phenoloxidase suppression in QX disease outbreaks among Sydney rock oysters (*Saccostrea glomerata*). Aquaculture. 2003;223:29–39.

25. Dove MC. Effects of estuarine acidification on survival and growth of the Sydney rock oyster *Saccostrea glomerata*. The University of New South Wales; 2003.

26. Green TJ, Jones BJ, Adlard RD, Barnes AC. Parasites, pathological conditions and mortality in QX-resistant and wild-caught Sydney rock oysters, *Saccostrea glomerata*. Aquaculture. 2008;280:35–8.

27. Kleeman SN, Adlard RD, Lester RJG. Detection of the initial infective stages of the protozoan parasite *Marteilia sydneyi* in *Saccostrea glomerata* and their development through to sporogenesis. International Journal for Parasitology. 2002;32:767–84.

28. Dang C, Lambert C, Soudant P, Delamare-Deboutteville J, Zhang MM, Chan J, et al. Immune parameters of QX-resistant and wild caught Saccostrea glomerata hemocytes in relation to Marteilia sydneyi infection. Fish & Shellfish Immunology. 2011;31:1034–40.

29. Dove MC, Nell JA, O’Connor WA. Evaluation of the progeny of the fourth-generation Sydney rock oyster Saccostrea glomerata (Gould, 1850) breeding lines for resistance to QX disease (*Marteilia sydneyi*) and winter mortality (*Bonamia roughleyi*). Aquaculture Research. 2013;44:1791–800.

30. Green TJ, Barnes AC. Inhibitor of REL/NF-КB is regulated in Sydney rock oysters in response to specific double-stranded RNA and *Vibrio alginolyticus*, but the major immune anti-oxidants EcSOD and Prx6 are non-inducible. Fish & Shellfish Immunology. 2009;27:260–5.

31. Morga B, Arzul I, Faury N, Segarra A, Chollet B, Renault T. Molecular responses of *Ostrea edulis* haemocytes to an *in vitro* infection with *Bonamia ostreae*. Developmental & Comparative Immunology. 2011;35:323–33.

32. Morga B, Renault T, Faury N, Arzul I. New insights in flat oyster *Ostrea edulis* resistance against the parasite *Bonamia ostreae*. Fish & Shellfish Immunology. 2012;32:958–68.

33. Kleeman SN, Adlard RD. Molecular detection of *Marteilia sydneyi*, pathogen of Sydney rock oysters. Diseases of Aquatic Organisms. 2000;40:137–46.

34. Adlard R, Worthington Wilmer J. Aquatic animal health subprogram: Validation of DNA-based (PCR) diagnostic tests suitable for use in surveillance programs for QX disease of Sydney rock oysters (Saccostrea glomerata) in Australia. Brisbane: Queensland Museum; 2003.

35. R Programming for Bioinformatics | Robert Gentleman | Taylor & Francis. https://www.taylorfrancis.com/books/mono/10.1201/9781420063684/programming-bioinformatics-robert-gentleman. Accessed 6 Jan 2025.

36. R: The R Project for Statistical Computing. https://www.r-project.org/. Accessed 6 Jan 2025.

37. Wickham H, Averick M, Bryan J, Chang W, McGowan LD, François R, et al. Welcome to the Tidyverse. Journal of Open Source Software. 2019;4:1686.

38. Firke S. janitor: Simple Tools for Examining and Cleaning Dirty Data. 2016;:2.2.0.

39. Nenadic N, Bar I, McDougall C. Code analysis for “Molecular responses to *Marteilia sydneyi* infection in the Sydney rock oyster *Saccostrea glomerata*.” 2024.

40. Song L, Florea L. Rcorrector: efficient and accurate error correction for Illumina RNA-seq reads. GigaScience. 2015;4:s13742–015-0089-y.

41. Rivera-Vicéns RE, Garcia-Escudero CA, Conci N, Eitel M, Wörheide G. TransPi—a comprehensive Transcriptome analysis PIpeline for de novo transcriptome assembly. Molecular Ecology Resources. 2022;22:2070–86.

42. Di Tommaso P, Chatzou M, Floden EW, Barja PP, Palumbo E, Notredame C. Nextflow enables reproducible computational workflows. Nat Biotechnol. 2017;35:316–9.

43. Chen S, Zhou Y, Chen Y, Gu J. fastp: an ultra-fast all-in-one FASTQ preprocessor. Bioinformatics. 2018;34:i884–90.

44. Quast C, Pruesse E, Yilmaz P, Gerken J, Schweer T, Yarza P, et al. The SILVA ribosomal RNA gene database project: improved data processing and web-based tools. Nucleic Acids Research. 2012;41:D590–6.

45. Hölzer M, Marz M. De novo transcriptome assembly: A comprehensive cross-species comparison of short-read RNA-Seq assemblers. GigaScience. 2019;8:giz039.

46. Wang S, Gribskov M. Comprehensive evaluation of de novo transcriptome assembly programs and their effects on differential gene expression analysis. Bioinformatics. 2017;33:327–33.

47. Manni M, Berkeley MR, Seppey M, Simão FA, Zdobnov EM. BUSCO Update: Novel and Streamlined Workflows along with Broader and Deeper Phylogenetic Coverage for Scoring of Eukaryotic, Prokaryotic, and Viral Genomes. Molecular Biology and Evolution. 2021;38:4647–54.

48. Smith-Unna R, Boursnell C, Patro R, Hibberd JM, Kelly S. TransRate: reference-free quality assessment of de novo transcriptome assemblies. Genome Res. 2016;26:1134–44.

49. Camacho C, Coulouris G, Avagyan V, Ma N, Papadopoulos J, Bealer K, et al. BLAST+: architecture and applications. BMC Bioinformatics. 2009;10:421.

50. Buchfink B, Xie C, Huson DH. Fast and sensitive protein alignment using DIAMOND. Nat Methods. 2015;12:59–60.

51. Petersen TN, Brunak S, von Heijne G, Nielsen H. SignalP 4.0: discriminating signal peptides from transmembrane regions. Nat Methods. 2011;8:785–6.

52. Krogh A, Larsson B, von Heijne G, Sonnhammer EL. Predicting transmembrane protein topology with a hidden Markov model: application to complete genomes. J Mol Biol. 2001;305:567–80.

53. Emanuelsson O, Brunak S, von Heijne G, Nielsen H. Locating proteins in the cell using TargetP, SignalP and related tools. Nat Protoc. 2007;2:953–71.

54. Jones P, Binns D, Chang H-Y, Fraser M, Li W, McAnulla C, et al. InterProScan 5: genome-scale protein function classification. Bioinformatics. 2014;30:1236–40.

55. The Galaxy Community. The Galaxy platform for accessible, reproducible, and collaborative data analyses: 2024 update. Nucleic Acids Research. 2024;52:W83–94.

56. Li H. Aligning sequence reads, clone sequences and assembly contigs with BWA-MEM. 2013.

57. Liao Y, Smyth GK, Shi W. featureCounts: an efficient general purpose program for assigning sequence reads to genomic features. Bioinformatics. 2014;30:923–30.

58. Ewels P, Magnusson M, Lundin S, Käller M. MultiQC: summarize analysis results for multiple tools and samples in a single report. Bioinformatics. 2016;32:3047–8.

59. Love MI, Huber W, Anders S. Moderated estimation of fold change and dispersion for RNA-seq data with DESeq2. Genome Biol. 2014;15:550.

60. Benjamini Y, Hochberg Y. Controlling the False Discovery Rate: A Practical and Powerful Approach to Multiple Testing. Journal of the Royal Statistical Society: Series B (Methodological). 1995;57:289–300.

61. Wu T, Hu E, Xu S, Chen M, Guo P, Dai Z, et al. clusterProfiler 4.0: A universal enrichment tool for interpreting omics data. The Innovation. 2021;2:100141.

62. Kolde R. pheatmap: Pretty Heatmaps. 2019.

63. Blighe K. kevinblighe/EnhancedVolcano. 2024.

64. He Y, Li X, Shi C, Li Y, Li Q, Liu S. Transcriptome profiling of the Pacific oyster (*Crassostrea gigas*) suggests distinct host immune strategy in response to *Vibrio alginolyticus* infection. Aquaculture. 2022;560:738563.

65. Gomes-dos-Santos A, Lopes-Lima M, Castro LFC, Froufe E. Molluscan genomics: the road so far and the way forward. Hydrobiologia. 2020;847:1705–26.

66. Liu F, Li Y, Yu H, Zhang L, Hu J, Bao Z, et al. MolluscDB: an integrated functional and evolutionary genomics database for the hyper-diverse animal phylum Mollusca. Nucleic Acids Research. 2021;49:D988–97.

67. Zannella C, Mosca F, Mariani F, Franci G, Folliero V, Galdiero M, et al. Microbial Diseases of Bivalve Mollusks: Infections, Immunology and Antimicrobial Defense. Marine Drugs. 2017;15:182.

68. Pfisterer K, Shaw LE, Symmank D, Weninger W. The Extracellular Matrix in Skin Inflammation and Infection. Front Cell Dev Biol. 2021;9.

69. Lian J, Lv L, Yao H, Lin Z, Dong Y. Genome-Wide Characterization and Analysis of Expression of the Histone Gene Family in Razor Clam, Sinonovacula constricta. Fishes. 2022;7:5.

70. Sayeed K, Parameswaran S, Beucler MJ, Edsall LE, VonHandorf A, Crowther A, et al. Human cytomegalovirus infection coopts chromatin organization to diminish TEAD1 transcription factor activity. eLife. 2024;13.

71. Wang K, del Castillo C, Corre E, Espinosa EP, Allam B. Clam focal and systemic immune responses to QPX infection revealed by RNA-seq technology. BMC Genomics. 2016;17.

72. Leite RB, Milan M, Coppe A, Bortoluzzi S, dos Anjos A, Reinhardt R, et al. mRNA-Seq and microarray development for the Grooved carpet shell clam, *Ruditapes decussatus*: a functional approach to unravel host -parasite interaction. BMC Genomics. 2013;14:741.

73. Kim JY, Kim YM, Cho SK, Choi KS, Cho M. Noble tandem-repeat galectin of Manila clam *Ruditapes philippinarum* is induced upon infection with the protozoan parasite *Perkinsus olseni*. Developmental & Comparative Immunology. 2008;32:1131–41.

74. Wang W, Song X, Wang L, Song L. Pathogen-derived carbohydrate recognition in molluscs immune defense. International Journal of Molecular Sciences. 2018;19:721.

75. Tasumi S, Vasta GR. A Galectin of Unique Domain Organization from Hemocytes of the Eastern Oyster (*Crassostrea virginica*) Is a Receptor for the Protistan Parasite *Perkinsus marinus*. The Journal of Immunology. 2007;179:3086–98.

76. Hannemann F, Bichet A, Ewen KM, Bernhardt R. Cytochrome P450 systems— biological variations of electron transport chains. Biochimica et Biophysica Acta (BBA) - General Subjects. 2007;1770:330–44.

77. Guengerich FP. Common and Uncommon Cytochrome P450 Reactions Related to Metabolism and Chemical Toxicity. Chem Res Toxicol. 2001;14:611–50.

78. Wei X-M, Lu M-Y, Duan G-F, Li H-Y, Liu J-S, Yang W-D. Responses of CYP450 in the mussel *Perna viridis* after short-term exposure to the DSP toxins-producing dinoflagellate *Prorocentrum lima*. Ecotoxicology and Environmental Safety. 2019;176:178–85.

79. Morga B, Renault T, Faury N, Chollet B, Arzul I. Cellular and molecular responses of haemocytes from *Ostrea edulis* during in vitro infection by the parasite *Bonamia ostreae*. International Journal for Parasitology. 2011;41:755–64.

80. Tanguy A, Guo X, Ford SE. Discovery of genes expressed in response to *Perkinsus marinus* challenge in Eastern (*Crassostrea virginica*) and Pacific (*C. gigas*) oysters. Gene. 2004;338:121–31.

81. Perrigault M, Tanguy A, Allam B. Identification and expression of differentially expressed genes in the hard clam, *Mercenaria mercenaria*, in response to quahog parasite unknown (QPX). BMC Genomics. 2009;10:377.

82. Green TJ, Dixon TJ, Devic E, Adlard RD, Barnes AC. Differential expression of genes encoding anti-oxidant enzymes in sydney rock oysters, *Saccostrea glomerata* (gould) selected for disease resistance. Fish & Shellfish Immunology. 2009;26:799–810.

83. He Y, Jouaux A, Ford SE, Lelong C, Sourdaine P, Mathieu M, et al. Transcriptome analysis reveals strong and complex antiviral response in a mollusc. Fish & Shellfish Immunology. 2015;46:131–44.

84. McDougall C. Reinvigorating the Queensland Oyster Industry. Brisbane, Queensland: Griffith University; 2020.

85. Green TJ, Barnes AC. Reduced Salinity, but Not Estuarine Acidification, Is a Cause of Immune-Suppression in the Sydney Rock Oyster *Saccostrea glomerata*. Marine Ecology Progress Series. 2010;402:161–70.

86. Manoharan RR, Prasad A, Pospíšil P, Kzhyshkowska J. ROS signaling in innate immunity via oxidative protein modifications. Front Immunol. 2024;15.

87. Yarosz EL, Chang C-H. The role of reactive oxygen species in regulating t cell-mediated immunity and disease. Immune Netw. 2018;18:e14.

88. Butt D, Shaddick K, Raftos D. The effect of low salinity on phenoloxidase activity in the Sydney rock oyster, *Saccostrea glomerata*. Aquaculture. 2006;251:159–66.

89. Gordy MA, Pila EA, Hanington PC. The role of fibrinogen-related proteins in the gastropod immune response. Fish & Shellfish Immunology. 2015;46:39–49.

90. Hanington PC, Zhang S-M. The Primary Role of Fibrinogen-Related Proteins in Invertebrates Is Defense, Not Coagulation. Journal of Innate Immunity. 2010;3:17–27.

91. Zhang H, Wang L, Song L, Song X, Wang B, Mu C, et al. A fibrinogen-related protein from bay scallop *Argopecten irradians* involved in innate immunity as pattern recognition receptor. Fish & Shellfish Immunology. 2009;26:56–64.

92. Huang B, Zhang L, Li L, Tang X, Zhang G. Highly diverse fibrinogen-related proteins in the Pacific oyster *Crassostrea gigas*. Fish & Shellfish Immunology. 2015;43:485–90.

93. Ho MKC, Su Y, Yeung WWS, Wong YH. Regulation of Transcription Factors by Heterotrimeric G Proteins. http://www.eurekaselect.com.

94. Sun L, Ye RD. Role of G protein-coupled receptors in inflammation. Acta Pharmacol Sin. 2012;33:342–50.

95. Wang K, Pales Espinosa E, Tanguy A, Allam B. Alterations of the immune transcriptome in resistant and susceptible hard clams (*Mercenaria mercenaria*) in response to Quahog Parasite Unknown (QPX) and temperature. Fish & Shellfish Immunology. 2016;49:163–76.

96. Martín-Gómez L, Villalba A, Abollo E. Identification and expression of immune genes in the flat oyster *Ostrea edulis* in response to bonamiosis. Gene. 2012;492:81–93.

97. Martín-Gómez L, Villalba A, Kerkhoven RH, Abollo E. Role of microRNAs in the immunity process of the flat oyster *Ostrea edulis* against bonamiosis. Infection, Genetics and Evolution. 2014;27:40–50.

98. Sokolova IM. Apoptosis in molluscan immune defense. Invertebrate Survival Journal. 2009;6:49–58.

99. Elmore S. Apoptosis: A Review of Programmed Cell Death. Toxicol Pathol. 2007;35:495–516.

100. Lau Y-T, Santos B, Barbosa M, Pales Espinosa E, Allam B. Regulation of apoptosis-related genes during interactions between oyster hemocytes and the alveolate parasite *Perkinsus marinus*. Fish & Shellfish Immunology. 2018;83:180–9.

101. Anderson RS. *Perkinsus marinus* secretory products modulate superoxide anion production by oyster (*Crassostrea virginica*) haemocytes. Fish & Shellfish Immunology. 1999;9:51–60.

102. Schott EJ, Pecher WT, Okafor F, Vasta GR. The protistan parasite *Perkinsus marinus* is resistant to selected reactive oxygen species. Experimental Parasitology. 2003;105:232–40.

103. Li Y, Zhang L, Qu T, Tang X, Li L, Zhang G. Conservation and divergence of mitochondrial apoptosis pathway in the Pacific oyster, Crassostrea gigas. Cell Death Dis. 2017;8:e2915–e2915.

104. Vaux DL, Cory S, Adams JM. Bcl-2 gene promotes haemopoietic cell survival and cooperates with c-myc to immortalize pre-B cells. Nature. 1988;335:440–2.

105. Joseph SJ, Fernández-Robledo JA, Gardner MJ, El-Sayed NM, Kuo C-H, Schott EJ, et al. The alveolate *Perkinsus marinus*: biological insights from est gene discovery. BMC Genomics. 2010;11.

106. Pales Espinosa E, Corre E, Allam B. Pallial mucus of the oyster *Crassostrea virginica* regulates the expression of putative virulence genes of its pathogen *Perkinsus marinus*. International Journal for Parasitology. 2014;44:305–17.

107. Canesi L, Barmo C, Fabbri R, Ciacci C, Vergani L, Roch P, et al. Effects of vibrio challenge on digestive gland biomarkers and antioxidant gene expression in *Mytilus galloprovincialis*. Comparative Biochemistry and Physiology Part C: Toxicology & Pharmacology. 2010;152:399–406.

108. Montagnani C, Avarre JC, de Lorgeril J, Quiquand M, Boulo V, Escoubas JM. First evidence of the activation of *Cg-timp*, an immune response component of pacific oysters, through a damage-associated molecular pattern pathway. Developmental & Comparative Immunology. 2007;31:1–11.

