## Supplementary figures and images for "Transcriptomic responses to *Marteilia sydneyi* infection in the Sydney rock oyster *Saccostrea glomerata*"

### Supplementary figure 1

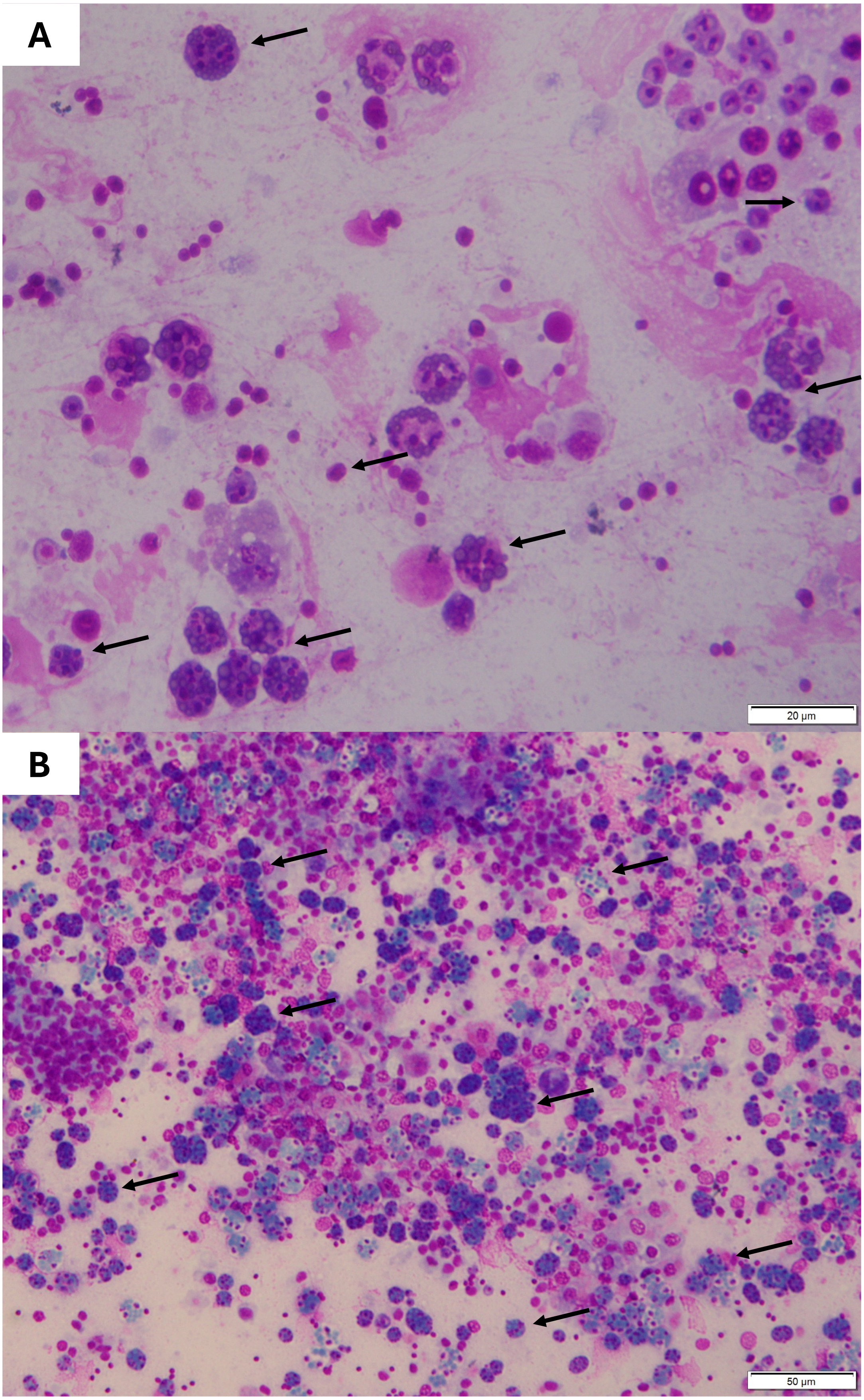

### Supplementary Figure 2

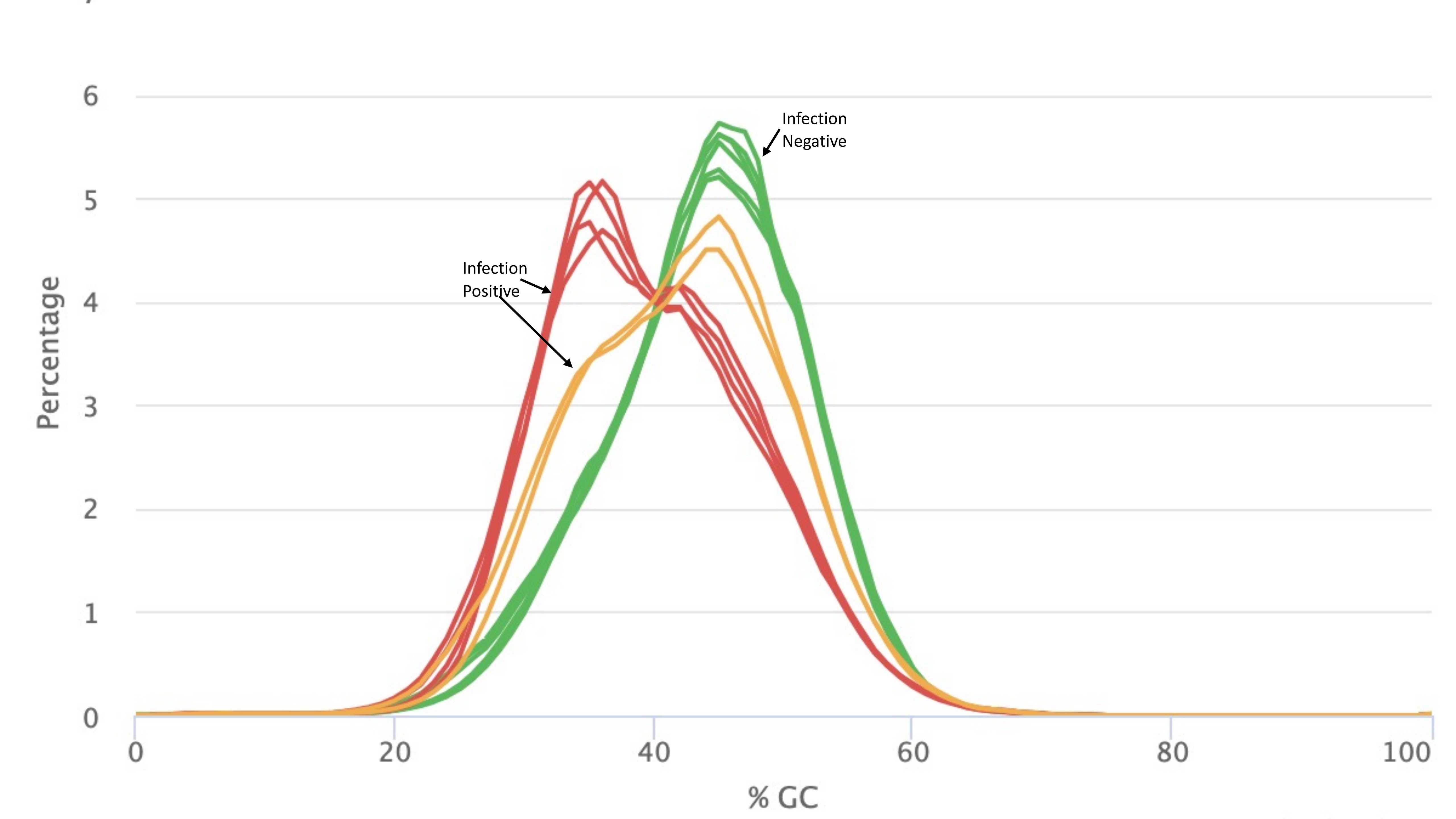
